# Osmotic Swelling Responses are Conserved Across Cartilaginous Tissues with Varied Sulfated-Glycosaminoglycan Contents

**DOI:** 10.1101/459115

**Authors:** Eva G. Baylon, Marc E. Levenston

**Author notes:** Corresponding Author: Building 520 Rm 225, Stanford University Stanford, CA 94305-4038, United States.

## Abstract

The interactions between the negatively charged sulfated glycosaminoglycan (sGAG) chains and the ionic interstitial fluid in articular cartilage and meniscal fibrocartilage give rise to an osmotic swelling stress that is critical for the load-bearing capability of both tissues. This osmotic swelling stress is altered when the sGAG content is changed, as during progression of degenerative joint disease; understanding the influence of sGAG concentration on the osmotic swelling stress of cartilage and meniscus is important to enhance our understanding of physiology and disease. This study compared the effect of altered osmotic environments on the confined compression swelling behavior of bovine tissues spanning a range of sGAG concentrations: juvenile articular cartilage, juvenile and adult meniscus, and juvenile cartilage degraded to reduce sGAG content. The transient response to changes in bath conditions was evaluated for explants assigned to one of three compressive offsets (5%, 10%, or 15% strain) and one of three bath conditions (0.1X, 1X, or 10X Phosphate Buffered Saline). Our results show that relative responses to alterations to the osmotic environment are consistent across tissue types, demonstrating that the role of sGAG in the swelling properties of the tissues tested is conserved, even when sGAG is present at low concentrations. Additionally, this study found unexpected correlations across tissue types between sGAG and collagen contents and between the aggregate modulus and both sGAG and collagen contents. These results suggest some conservation of composition-function relationships across a range of tissue types.

## Introduction

Articular cartilage and meniscal fibrocartilage transfer loads and provide lubrication and stability in the knee joint.^1^ Both cartilage and meniscus are comprised primarily of water (68-85%, 60-70%, respectively), collagen (10-20% type II in cartilage, 15-25% type I in meniscus), and proteoglycans (5-10%, 1-2% respectively), predominantly aggrecan.^2^ Structurally, collagen is mainly responsible for the tensile stiffness of cartilage and meniscus. The equilibrium compressive stiffness of these tissues arises in part from an osmotic swelling stress due to interactions between the ionic interstitial fluid and the negatively charged sulfated glycosaminoglycan (sGAG) chains that are covalently attached to the proteoglycan core protein. This osmotic swelling stress helps to maintain the structure and mechanical function of both cartilage and meniscus, both directly by resisting tissue compression and indirectly by sustaining tension in the collagen network that contributes to tensile and shear stiffness.^3^ Intra‐ and inter-tissue differences in tissue composition produce variations in the baseline swelling stress, and loss of GAGs during degeneration can substantially reduce the swelling stress and consequently impair mechanical function. Additionally, local tissue compression can force fluid out of the tissue, effectively increasing the local sGAG concentration and the resulting swelling stress.

Typically, experiments to assess the proteoglycan-associated osmotic swelling stress involve changing the osmotic environment by altering the salt concentration of the specimen bath. This type of alteration, although not observed under physiological conditions, is an important laboratory tool for probing these types of charge interactions and understanding the influence of swelling stress in physiology and pathology. Previous studies have established that decreases in the sGAG concentration, such as those that occur with osteoarthritis, result in a reduction of the osmotic swelling stress in both cartilage and meniscus, while decreases to the bath salinity result in increases to the osmotic swelling stress (and conversely for increases in these quantities).^4–10^ Nonetheless, the effects of the interaction between both sGAG concentration and changes to bath salinity on the transient osmotic swelling stress of cartilaginous tissues remains unclear. The goal of this study was to assess the transient osmotic swelling response to altered osmotic environments in tissues with a range of sGAG concentrations. We hypothesized that, while relative changes to the osmotic swelling stress would be similar regardless of sGAG concentration, the transient behavior would differ across tissue types.

## Materials and Methods

### Materials

Juvenile (approximately 2 week old) bovine stifles were from Research 87, Inc. (Boylston, MA). Adult bovine stifles (approximately 2 years old) were acquired from The Local Butcher Shop (Berkeley, CA). Biopsy punches were from Integra Miltex (York, PA). Protease Inhibitor Cocktail Set I and Phosphate Buffered Saline (PBS) tablets (140mM NaCl, 10mM phosphate buffer, 3mM KCl reconstituted in 1L of deionized H2O) were from Calbiochem (San Diego, CA). Chondroitinase-ABC (Ch-ABC), 1,9-dimethyl-methylene blue (DMMB) dye, shark chondroitin sulfate, and trans-4-hydroxy-_L_-proline were from Sigma Aldrich (St. Louis, MO). Proteinase K was from Fisher Scientific (Hampton, NH).

### Sample preparation

Juvenile cartilage and meniscus samples were harvested from 6 juvenile bovine stifles and adult meniscus samples were obtained from 3 adult bovine stifles. Full thickness cores of cartilage from the femoral condyles and both medial and lateral menisci were removed with a 6mm biopsy punch. Samples were trimmed to a final thickness of 1mm using a sliding microtome equipped with a freezing stage, discarding the superficial and deep zones, and stored until the day of testing at −20°C in 1X PBS supplemented with protease inhibitors. A subset of cartilage cores were treated with Ch-ABC to enzymatically reduce the sGAG concentration using a protocol determined in preliminary studies to produce a final sGAG concentration similar to that of adult meniscus (see supplemental data, Fig. S1). Cartilage cores trimmed to 2mm thickness were first equilibrated in 1X PBS supplemented with 0.3ml of 1U/ml of Ch-ABC overnight at 4°C to ensure homogenous distribution of the enzyme throughout the explant (Ch-ABC has minimal activity at 4°C^11^), and subsequently incubated at 37°C for 42hrs. After Ch-ABC incubation, cartilage explants were washed in 1X PBS three times for 30 minutes at room temperature to remove any excess enzyme. Explants were frozen and the top and bottom layers were removed with a microtome to leave a 1mm-thick sample. On the day of testing, explants were thawed and biopsy-punched to a final diameter of 4mm.

### Confined compression testing

All mechanical testing was performed at room temperature. Sample wet masses and thicknesses were recorded prior to mechanical testing. Samples were placed in a custom-made aluminum rig consisting of a 4mm-diameter chamber with a stainless steel porous filter lining the bottom of the chamber that allowed fluid to be exuded out of the tissue due to compression. Fluid was recirculated into the bath with a peristaltic pump throughout the test. Samples were subjected to a confined compression regime in this rig in an Instron 5848 Microtester (Instron, Norwood, MA, USA) using either a 1N (FUTEK Advanced Sensor Technology, Inc., Irvine, CA) or 10N (Interface, Inc., Scottsdale, AZ) load cell, with compression applied to the top of the sample via a flat 4mm-diameter aluminum rod.

Juvenile meniscus, juvenile cartilage, degraded juvenile cartilage, and adult meniscus samples were assigned to one of three bath tonicities - 0.1X PBS (hypotonic), 1X PBS (isotonic), or 10X PBS (hypertonic) - and one of three compressive offsets - 5%, 10%, or 15% strain - for a total of nine experimental groups per tissue type (Fig. 1; n=5/group/tissue). After applying a tare load of - 0.02N and adding 1X PBS to the chamber, the assigned compressive offset was applied at a rate of 0.001mm/s and the samples were allowed to stress-relax for 45 minutes. Upon completion of the relaxation period, the PBS bath was changed to the assigned tonicity group and the samples were allowed to re-equilibrate for an additional 90 minutes. All samples were then unloaded at a rate of 0.01mm/s, removed from the confined compression rig and allowed to re-equilibrate in 1X PBS for an hour. The final wet mass was recorded before storing samples at −20°C for biochemical analysis.

**Figure 1:**
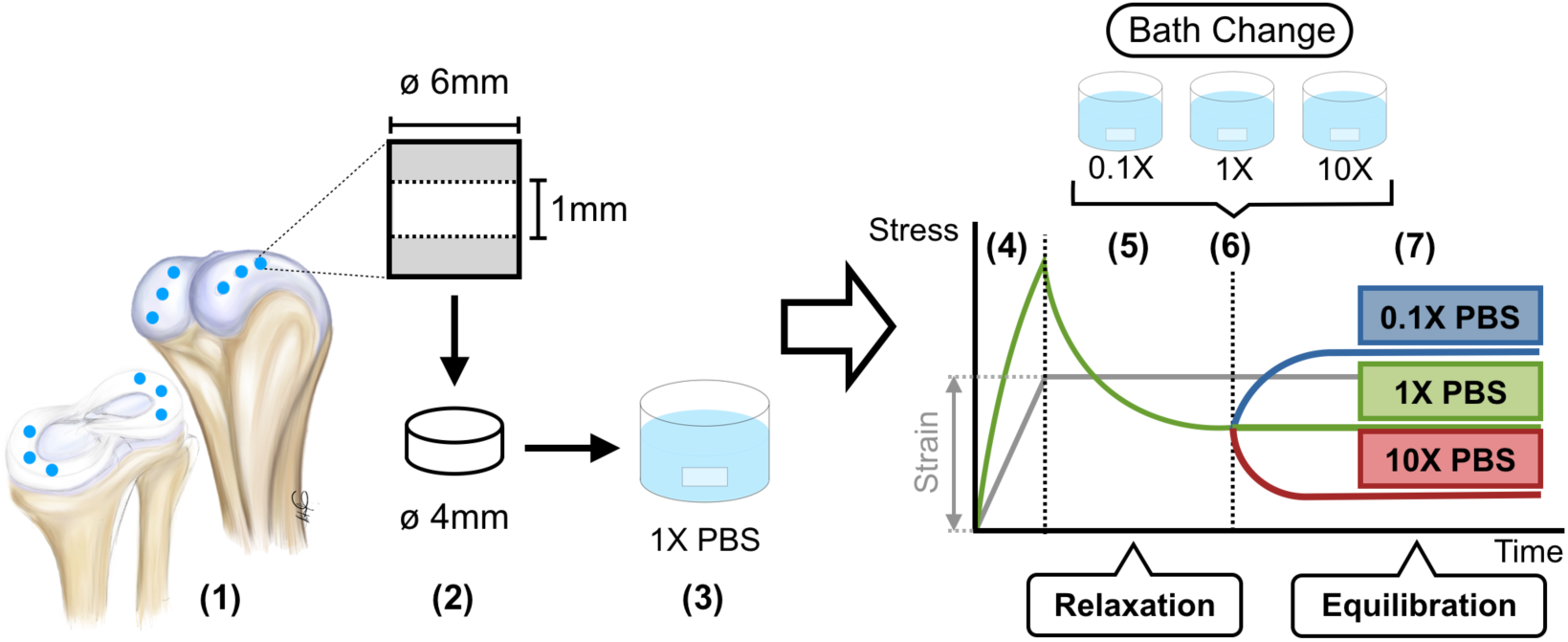
Osmotic swelling experiment set up. (1) Full thickness (6mm diameter) meniscus and cartilage cores were harvested from juvenile and adult bovine stifles and (2) trimmed to a final thickness of 1mm and diameter of 4mm. A subset of juvenile cartilage samples were enzymatically treated with chondroitinase ABC to reduce the sGAG content, as described in the text. (3) All samples were equilibrated in 1X PBS prior to mechanical testing. (4) 5%, 10%, or 15% compressive strain was applied on the sample in a custom made confined compression rig at 0.001mm/s in 1X PBS. (5) Samples were allowed to stress-relax for 45 minutes. (6) Bath was changed to 0.1X PBS (hypotonic), 1X PBS (isotonic control), or 10X PBS (hypertonic) and recirculated with a pump throughout the test. (7) Samples were allowed to re-equilibrate for an additional 90 minutes.

### Data Analysis

A linear biphasic model^12,13^ was fitted to the loading and initial 45min relaxation data using a least squares approach in MATLAB (version R2014b, MathWorks, Natick, MA) to estimate the aggregate modulus (H_A_), which describes the equilibrium relationship between stress and strain in confined compression. The change in swelling stress was quantitatively analyzed with the swelling stress ratio (equation 1), the ratio between the equilibrium swelling stress after changing the bath to the equilibrium swelling stress before changing the bath.

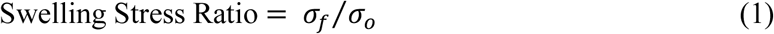

The second transient, related to the re-equilibration after changing the bath, was characterized with a decaying mono-exponential equation to obtain the osmotic equilibrium time constant τ (equation 2)

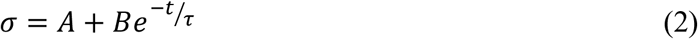

### Biochemical analysis

Explants were lyophilized, weighed dry, and digested in 0.5ml of 2mg/ml Proteinase K in 100mM ammonium acetate at 60°C overnight following mechanical testing. Water content was estimated using the dry and wet masses of the samples. The sGAG content was analyzed using the 1,9-dymethylmethylene blue (DMMB) spectrophotometric assay at 595nm-525nm, pH=3.0^14^, using chondroitin sulfate C standards. To estimate the collagen content, the hydroxyproline content was measured via the p-dimethylaminobenzaldehyde spectrophotometric assay at 557nm with trans-4-hydroxy-_L_-proline standards and converted to collagen content using a collagen to hydroxyproline mass ratio of 8.^15^

### Statistical analysis

Swelling stress ratio and equilibrium time constant data were analyzed with multi-factor general linear models (GLMs) with tissue type, compressive offset, and PBS group as fixed factors. Bonferroni’s post-hoc test was used for comparisons for main and interaction effects, with significance set at p<0.05. All data were processed with Box-Cox transformation to improve normality. The corrected Akaike information criterion (AICc)^16^ was minimized to determine best models through backwards selection. Correlations between mechanical parameters and tissue composition were done using Spearman’s correlation analysis. Nonlinear regression analyses of aggregate modulus vs collagen/dry mass and collagen/dry mass vs sGAG/dry mass data sets were used to determine power law functions while a linear regression was used for the aggregate modulus vs sGAG/dry mass data set. All statistical analyses were performed using Minitab (version 17, Minitab, Inc., State College, PA). Results are presented as mean±SEM.

## Results

### Biochemical composition

Tissue composition is summarized in Table 1. Water content did not significantly differ among meniscus groups or among cartilage groups, but it was significantly lower overall for the meniscus groups compared to the cartilage groups (Fig. 2A). The lowest sGAG concentration was measured for juvenile meniscus while the highest concentration was observed for juvenile cartilage. There was no significant difference between the sGAG concentrations of adult meniscus and degraded cartilage, and both groups had significantly higher sGAG concentrations than juvenile meniscus and significantly lower concentrations than juvenile cartilage (Fig. 2B). The collagen concentration was significantly different amongst all groups, with juvenile meniscus having the highest collagen/wet mass and juvenile cartilage having the lowest (Fig. 2C).

**Table 1.** Biochemical composition of all tissues tested.

| Tissue Type | Water (% w/w) | sGAG/wet mass ($\mu\text{g}/\text{mg}$) | Collagen/wet mass ($\mu\text{g}/\text{mg}$) |
| --- | --- | --- | --- |
| Juvenile Meniscus | 71.45±0.45 | 3.50 ±0.38 | 334.75±7.67 |
| Adult Meniscus | 72.05±0.43 | 11.23±0.89 | 229.44±6.64 |
| Degraded Cartilage | 78.83±2.17 | 16.64±3.18 | 154.68±4.54 |
| Juvenile Cartilage | 79.62±0.39 | 67.71±2.69 | 131.00±4.57 |

**Figure 2:**
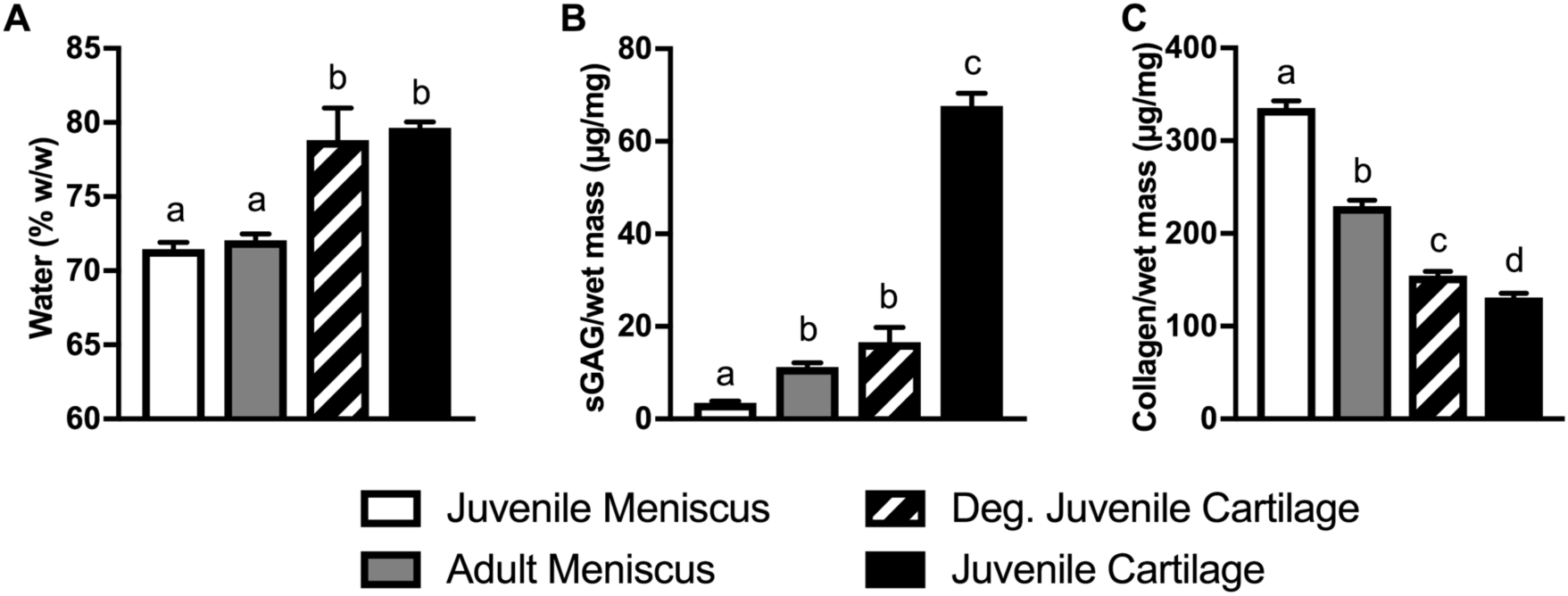
Biochemical composition of tissue types tested (n=45). (A) Water content (% wet weight), (B) sGAG/wet mass (μg/mg) and (C) collagen/wet mass (μg/mg). Columns that do not share a letter are significantly different (p<0.05).

### Aggregate modulus

The aggregate modulus (Fig. 3) was lowest for juvenile meniscus (29.81±1.72kPa) and highest for articular cartilage (378.92±23.34kPa), with no significant difference between adult meniscus (55.47±2.82kPa) and degraded cartilage (72.4±8.44kPa).

**Figure 3:**
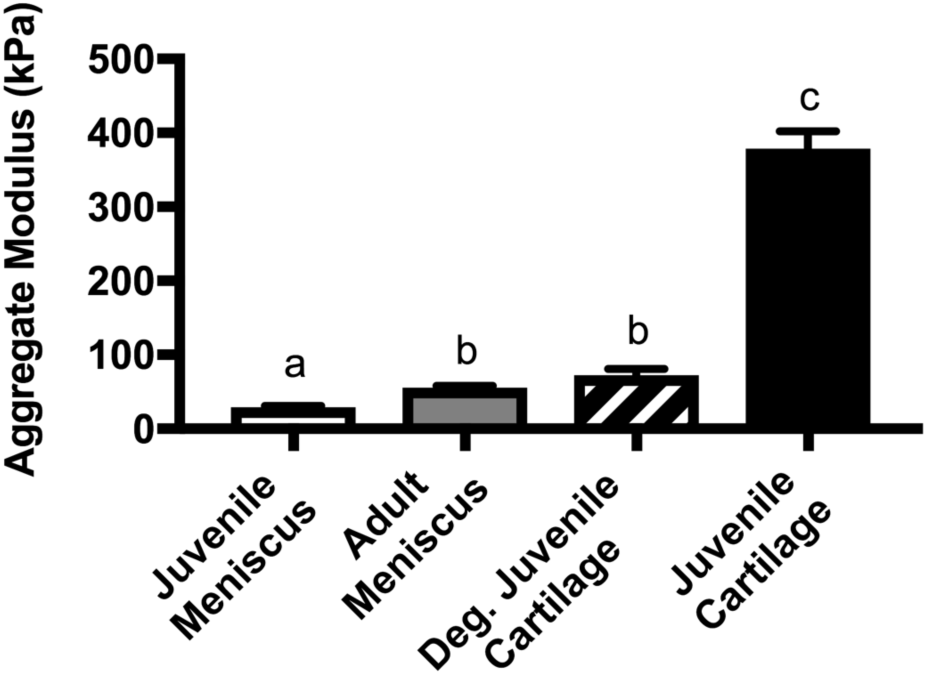
Comparison of aggregate modulus for all tissue types (n=45). The aggregate modulus was estimated by fitting a linear biphasic model to the initial portion (ramp and 45 minute relaxation) of the experimental data. The aggregate modulus was significantly lower for juvenile meniscus than for other tissue groups while that of juvenile cartilage was significantly higher than for other groups. Columns that do not share a letter are significantly different (p<0.05).

### Osmotic swelling stress ratio

Compressive offset (5%, 10%, 15%) did not significantly influence the swelling stress ratio. To simplify presentation, results are presented as pooled across compressive offset groups for a given tissue type and bath condition (Table 2; n=15/group). The swelling stress ratio patterns were consistent for all tissue types and as expected (Fig. 4), with a stress ratio greater than one for the 0.1X PBS groups (indicating swelling), approximately one for the 1X PBS groups (indicating no change), and less than one for the 10X PBS groups (indicating deswelling). The stress ratio for 10X PBS was significantly lower than for 1X PBS for every tissue group except degraded cartilage, while the stress ratio for 0.1X PBS was significantly greater than for 1X PBS for juvenile cartilage and adult meniscus. No correlations were observed between swelling stress ratios and biochemical composition (Fig. S2). A comparison of native tissues shows no significant differences between the swelling stress ratios at 1X PBS or 10X PBS among tissue types. For the 0.1X PBS group, the swelling stress ratio for juvenile meniscus was significantly lower than for adult meniscus or juvenile cartilage (Fig. 5A).

**Table 2.** Swelling stress ratios for all PBS equilibration groups.

| Tissue Type | Stress Ratio ($\sigma_f/\sigma_o$) | | |
| --- | --- | --- | --- |
|  | 0.1X PBS | 1X PBS | 10X PBS |
| Juvenile Meniscus | 1.231±0.191 | 0.863±0.033 | 0.549±0.109 |
| Adult Meniscus | 1.714±0.078 | 0.905±0.012 | 0.448±0.056 |
| Degraded Cartilage | 1.217±0.083 | 0.955±0.014 | 0.752±0.061 |
| Juvenile Cartilage | 1.869±0.177 | 1.028±0.017 | 0.356±0.019 |

**Figure 4:**
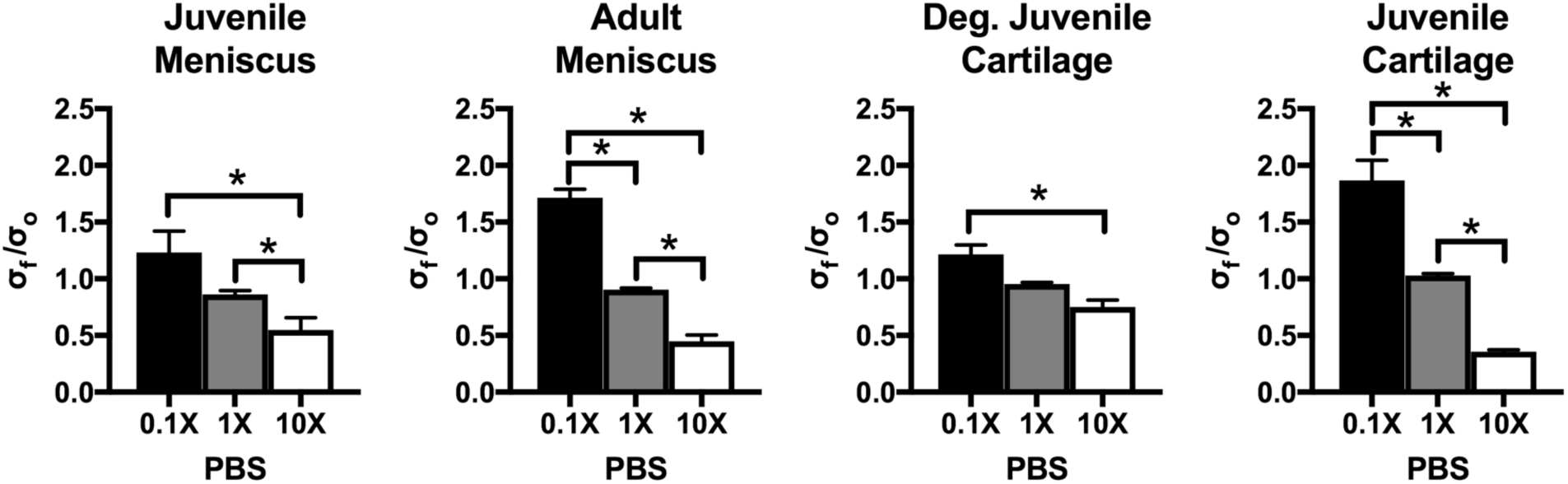
Swelling stress ratios for all tissue types tested, pooled across compressive offset groups. Swelling stress ratios were consistent across tissue types, with a value greater than 1 for groups equilibrated in 0.1XPBS, approximately 1 for groups equilibrated in 1X PBS, and less than 1 for groups equilibrated in 10X PBS.* indicates significant differences between groups (p<0.05).

**Figure 5:**
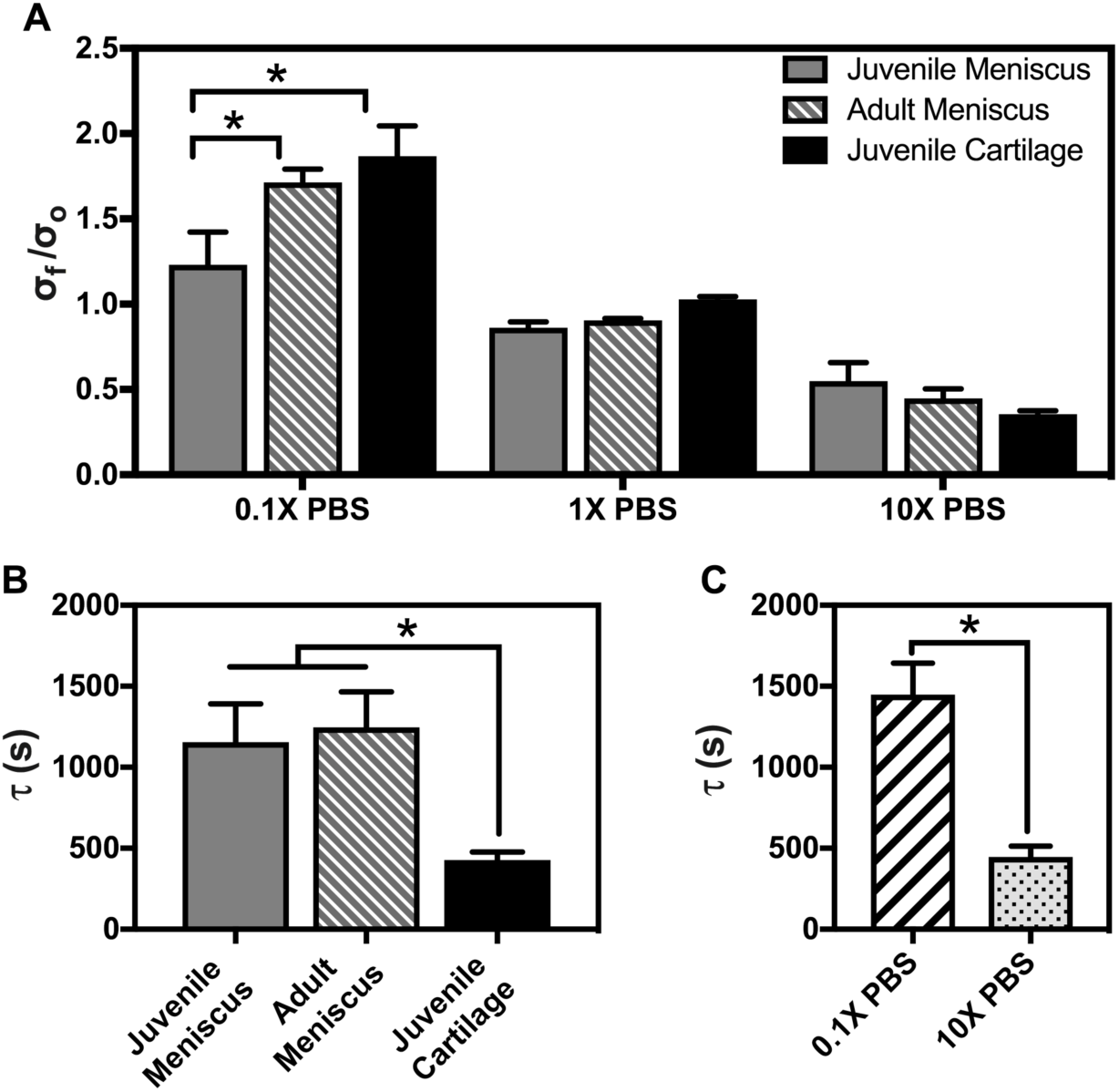
Comparison of swelling stress ratios and equilibrium time constants among native tissue types, pooled across compressive offsets. (A) The swelling stress ratio was only significantly different between juvenile meniscus and adult meniscus and juvenile cartilage for the groups equilibrated in 0.1X PBS (n=15/group). (B) The equilibrium time constant was significantly lower for juvenile cartilage than for juvenile and adult meniscus (n=45/group). (C) The equilibrium time constant across tissue types (n=45/group) shows that swelling (0.1X PBS) takes significantly longer than deswelling (10X PBS). * indicate differences between groups are significant (p<0.05).

### Osmotic equilibrium time constant

Consistent with the swelling stress ratio results, compressive offset did not significantly affect the osmotic equilibrium time constant, and Fig. 6 presents these data pooled across compressive offset groups for each PBS concentration and tissue type (n=15/group). For all native tissues (all tissues except degraded cartilage), equilibration in 0.1X PBS (swelling) took significantly longer than equilibration in 10X PBS (deswelling). These results are summarized in Table 3. No correlations were found between equilibrium time constants and biochemical composition (Fig. S2). Focusing only on the native tissues, the osmotic equilibrium time constant was significantly lower for juvenile cartilage (427.8±48.3s) when compared to juvenile (1155.9±235.1s) and adult (1246.4±219.1s) meniscus tissues (Fig. 5B) across PBS and compressive offset groups. The osmotic equilibrium time constant was significantly greater for swelling (1449±193.6s) than deswelling (447.0±66.5s) across tissue types and compressive offset groups (Fig. 5C).

**Table 3.** Osmotic equilibrium time constant results summary.

| Tissue Type | Osmotic Equilibrium Time Constant (s) |  |
| --- | --- | --- |
|  | 0.1X PBS | 10X PBS |
| Juvenile Meniscus | $1789 \pm 393.8$ | $522.6 \pm 129.3$ |
| Adult Meniscus | $1889 \pm 341.9$ | $603 \pm 137.4$ |
| Degraded Cartilage | $739.9 \pm 138.6$ | $847.8 \pm 267.2$ |
| Juvenile Cartilage | $644.7 \pm 44.8$ | $225.5 \pm 35.2$ |

**Figure 6:**
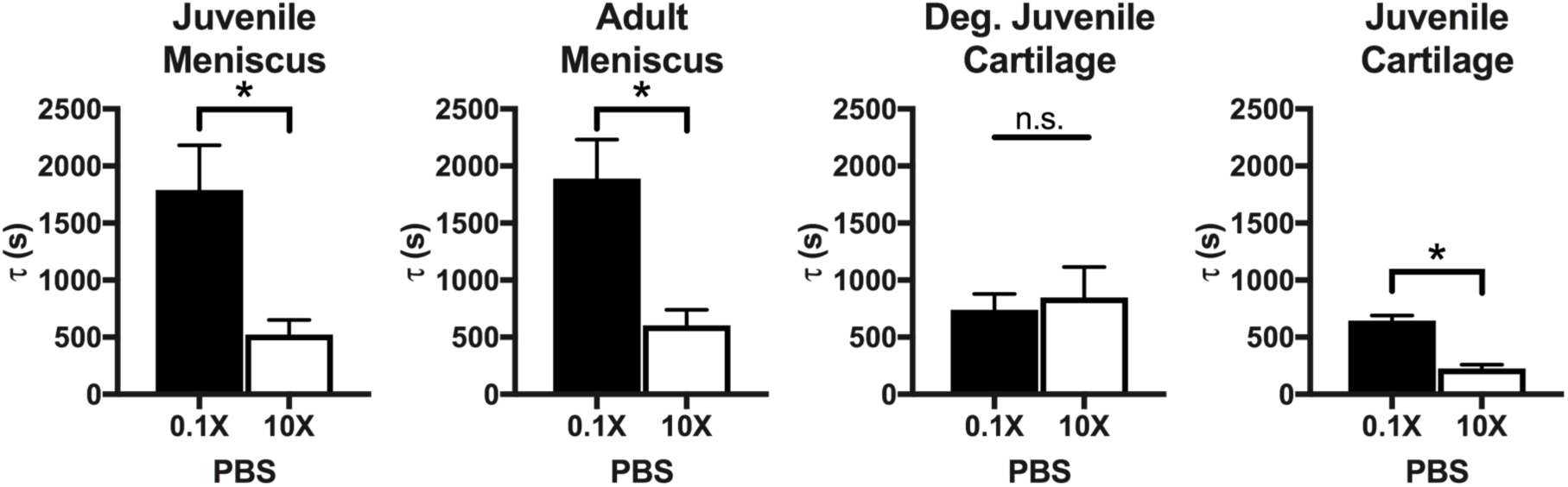
Comparison of osmotic equilibrium time constants between all tissue types tested across compressive offset groups. The equilibrium time constant for swelling (0.1X PBS) was significantly longer than for deswelling (10X PBS) for all tissue types except degraded cartilage. * indicate differences between groups are significant (p<0.05).

### Correlation of mechanical parameters to tissue composition

Spearman’s correlation analysis of native tissues indicates a strong negative correlation between the aggregate modulus and collagen/dry mass (rho=-0.778, p<0.001) and a strong positive correlation between the aggregate modulus and sGAG/dry mass (rho=0.850, p<0.001) (Fig. 7). The aggregate modulus-collagen/dry mass relationship was best described with a power fit (HA=45875[col/dm]’^−3.117^, R^2^=0.599), while the aggregate modulus-sGAG/dry mass relationship was characterized with a linear regression (H_A_=881396[sGAG/dm]+39364, R^2^=0.592). Interestingly, across native tissues the collagen and sGAG normalized to dry mass exhibited a strong inverse correlation (rho=-0.803, p<0.001; Fig. 8). Similar trends were observed when biochemical composition was normalized by wet mass or water content and those results are included as supplemental data (Fig. S3).

**Figure 7:**
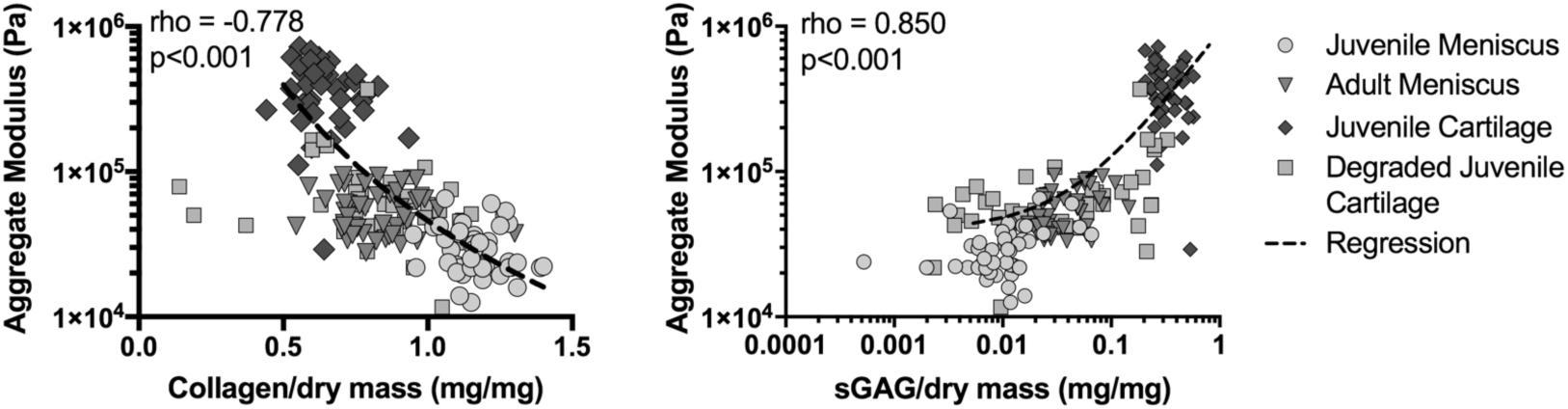
Correlation analysis between aggregate modulus (H_A_) and (A) Collagen/dry mass (regression equation: H_A_=45875[col/dm]^−3.117^, R^2^=0.599) and (B) sGAG/dry mass (regression equation: H_A_=881396[sGAG/dm]+39364, R^2^=0.592) in native tissues along with Spearman’s correlation analyses. A negative correlation was found between aggregate modulus and collagen/dry mass while a positive correlation was found between aggregate modulus and sGAG/dry mass across tissue types. Note that aggregate modulus vs collagen/dry mass is a semi-log plot while aggregate modulus vs sGAG/dry mass is a log-log plot. Plots include degraded cartilage data for comparison (excluded from analysis).

**Figure 8:**
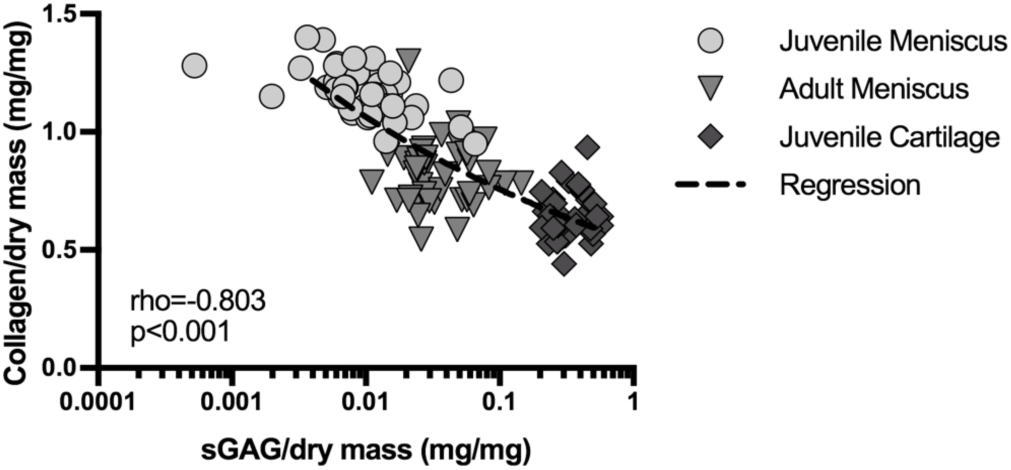
Correlation analysis between collagen/dry mass and sGAG/dry mass (regression equation: [Col/dm]=0.5385[sGAG/dm]^−0.148^, R^2^=0.657) along with Spearman’s correlation analysis. A negative correlation was found between collagen/dry mass and sGAG/dry mass across tissues types. Note that sGAG/dry mass is on a log scale.

## Discussion

Biochemical composition of bovine tissue explants was consistent with previous reports^5,17,18^, with sGAG content lowest for juvenile meniscus and highest for juvenile cartilage. As expected from our preliminary experiments to optimize Ch-ABC concentration and incubation times, enzymatic treatment of juvenile cartilage explants resulted in similar sGAG tissue concentrations to that of adult meniscus explants. Collagen content was significantly different among all groups, including degraded and juvenile cartilage. Differences in collagen concentration in degraded and juvenile cartilage groups might be a result of the overall reduction in the mass of degraded cartilage as sGAGs are depleted from the tissue (a reduction in mass would lead to a higher collagen/wet mass). This is also supported by previous studies that have used Ch-ABC to enzymatically degrade the GAGs in the extracellular matrix while keeping the collagen network intact.^19,20^

Swelling stress ratios for all tissue types varied as expected, with a stress ratio greater than 1 (indicative of swelling) for equilibration in 0.1X PBS, approximately 1 for equilibration in 1X PBS (control), and less than 1 (indicative of deswelling) for equilibration in 10X PBS. Although the actual values of the osmotic swelling stresses were orders of magnitude different between meniscus and cartilage explants, the relative changes in the osmotic swelling stress were similar across tissue types, suggesting that the role of sGAG in cartilaginous tissue swelling properties is conserved even at low concentrations.

Compressive offset did not have an effect on the swelling stress ratio or the equilibrium time constant results. Previous studies have shown that dynamic, shear, and equilibrium moduli in unconfined compression increase with compressive offset.^5^ Although confined compression effectively changes the concentration of sGAG and collagen by reducing the total fluid volume as fluid is exuded from the sample, this effect is negligible for the swelling and transport properties of cartilage and meniscus compared to the large effects of altering the osmotic environment by changing the ionic concentration.

Equilibrium time constants for meniscus specimens were longer than for cartilage specimens. This equilibration is governed by both the apparent diffusivity – which affects the transport of ions – and the permeability – which influences the transient fluid redistribution. Both of these intrinsic parameters are related to structure and sGAG-ion interactions, with longer equilibration times observed in tissues with higher densities and lower sGAG concentrations, consistent with the results from this study.

For all native tissues, swelling took significantly longer than deswelling, regardless of compressive offset. In confined compression, the volume of the explants remains constant throughout the test and changes to the osmotic environment only lead to local changes as a result of transient fluid redistribution within the sample that eventually returns to its baseline distribution. Equilibration in a hypotonic bath increases the Debye length, the effective distance over which the negative fixed charges perturb the local ion distribution^21^. As a result, ions within the extracellular matrix will interact more with these negative charges, slowing down their transport and creating a more tortuous path that leads to longer equilibration times.

Previous studies have widely established the role of sGAG in the mechanical properties of cartilage and meniscus in compression.^22^ As a result, the correlation between sGAG concentration and the aggregate modulus was expected. Nonetheless, the fact that this correlation holds across tissue types suggests that there exists a similar association even for the lowest sGAG concentration tissue (juvenile meniscus). The strong association observed between collagen and sGAG concentrations across tissue types suggests that the relationship between sGAG and collagen might be preserved for the tissues examined here. Furthermore, this trend seems to hold for other types of tissues outside the realm of the sGAG concentration spectrum tested, particularly for tendon (with collagen/dry mass values in the order of 100μg/mg and sGAG/dry mass of approximately 1μg/mg for flexor bovine tendons in 6mo. old calves).^23^

Overall, these results suggest that both sGAG and collagen, and the interaction between these two tissue components, are fundamental for maintaining optimal tissue compressive behaviors. Future studies should consider conditions under which the relationship between sGAG and collagen might break down to mimic scenarios typically observed with degenerative joint disease. Additionally, a computational transport model could be a better tool to describe the response to alterations to the osmotic environment, lending more direct insights into the fundamental physical phenomena at play.

## Acknowledgements

The studies presented here were supported by a National Science Foundation Graduate Research Fellowship, a Stanford Bio-X Fellowship, an Achievement Rewards for College Scientists Fellowship, and a Sloan PhD Fellowship (EGB). The authors would like to acknowledge Heidi G. Poppe and Francisco Lopez for assisting with data collection. The content is solely the responsibility of the authors and does not represent the official views of the funders.

Author Contribution Statement
All authors contributed to study design. Studies were performed by EGB and analyzed by EGB and MEL. All authors contributed to data interpretation and writing the manuscript. All authors reviewed and approve of the final manuscript.

## References

1. McDevitt CA, Webber RJ. 1990. The Ultrastructure and Biochemistry of Meniscal Cartilage. Clin. Orthop. Relat. Res. (252):8–18.

2. Mow VC, Huiskes R. 2005. Basic Orthopaedic Biomechanics & Mechano-biology. Philadelphia: Williams & Wilkins.

3. Fithian DC, Kelly MA, Mow VC. 1990. Material Properties and Structure-Function Relationships in the Menisci. Clin. Orthop. Relat. Res. (252): 19–31.

4. Eisenberg SR, Grodzinsky AJ. 1985. Swelling of Articular Cartilage and Other Connective Tissues: Electromechanochemical Forces. J. Orthop. Res. 3(2): 148–159.

5. Nguyen AM, Levenston ME. 2012. Comparison of Osmotic Swelling Influences on Meniscal Fibrocartilage and Articular Cartilage Tissue Mechanics in Compression and Shear. J. Orthop. Res. 30(1):95–102.

6. Sun DD, Guo XE, Likhitpanichkul M, et al. 2004. The Influence of the Fixed Negative Charges on Mechanical and Electrical Behaviors of Articular Cartilage Under Unconfined Compression. J. Biomech. Eng. 126(1):6.

7. Korhonen RK, Jurvelin JS. 2010. Compressive and tensile properties of articular cartilage in axial loading are modulated differently by osmotic environment. Med. Eng. Phys. 32:155–160.

8. Bursac P, Arnoczky S, York a. 2009. Dynamic compressive behavior of human meniscus correlates with its extra-cellular matrix composition. Biorheology 46(3):227–37.

9. Andrews SHJ, Rattner JB, Shrive NG, Ronsky JL. 2015. Swelling significantly affects the material properties of the menisci in compression. J. Biomech. 48(8):1485–1489.

10. Wilson CG, Vanderploeg EJ, Zuo F, et al. 2009. Aggrecanolysis and in vitro matrix degradation in the immature bovine meniscus: mechanisms and functional implications. Arthritis Res. Ther. 11(6):R173.

11. Shamsi M, Akram Shirdel S, Jafarian V, et al. 2016. Optimization of conformational stability and catalytic efficiency in chondroitinase ABC I by protein engineering methods. Eng. Life Sci 16:690–696.

12. Mow VC, Kuei SC, Lai WM, Armstrong CG. 1980. Biphasic creep and stress relaxation of articular cartilage in compression: Theory and experiments. J. Biomech. Eng. 102(1):73–84.

13. Bursać PM, Obitz TW, Eisenberg SR, Stamenović D. 1999. Confined and unconfined stress relaxation of cartilage: Appropriateness of a transversely isotropic analysis. J.. 32(10): 1125–1130.

14. Zheng CH, Levenston ME. 2015. Fact versus artifact: avoiding erroneous estimates of sulfated glycosaminoglycan content using the dimethylmethylene blue colorimetric assay for tissue-engineered constructs. Eur. Cell. Mater. 29:224–36; discussion 236.

15. Woessner JF. 1961. The determination of hydroxyproline in tissue and protein samples containing small proportions of this imino acid. Arch. Biochem. Biophys. 93(2):440–447.

16. Burnham KP, Anderson DR. 2004. Multimodel inference: Understanding AIC and BIC in model selection. Sociol. Methods Res. 33(2):261–304.

17. Ling CH-Y, Lai JH, Wong IJ, Levenston ME. 2016. Bovine Meniscal Tissue Exhibits Age-and Interleukin-1 Dose-Dependent Degradation Patterns and Composition-Function Relationships. J Orthop Res 34:801–811.

18. Fermor HL, McLure SWD, Taylor SD, et al. 2015. Biological, biochemical and biomechanical characterisation of articular cartilage from the porcine, bovine and ovine hip and knee. Biomed. Mater. Eng. 25:381–395.

19. Asanbaeva A, Masuda K, Thonar EJ, et al. 2007. Mechanisms of cartilage growth: Modulation of balance between proteoglycan and collagen in vitro using chondroitinase ABC. Arthritis Rheum. 56(1): 188–198.

20. Bautista CA, Park HJ, Mazur CM, et al. 2016. Effects of Chondroitinase ABC-Mediated Proteoglycan Digestion on Decellularization and Recellularization of Articular Cartilage. PLoS One 11(7):e0158976.

21. Mattern KJ, Nakornchai C, Deen WM. 2008. Darcy Permeability of Agarose-Glycosaminoglycan Gels Analyzed Using Fiber-Mixture and Donnan Models. Biophys. J. 95(2):648–656.

22. Maroudas A. 1979. Physico-Chemical Properties of Articular Cartilage. In: Freeman MAS, editor. Adult Articular Cartilage. Turnbridge Wells: Pitman Medical. p 215–290.

23. Evanko SP, Vogel KG. 1990. Ultrastructure and Proteoglycan Composition in the Developing Fibrocartilaginous Region of Bovine Tendon. Matrix Collagen Relat. Res. 10:420–436.

